# Single cell RNA-sequencing reveals an association between testosterone treatment and reduced hormone signaling in the human mammary gland

**DOI:** 10.64898/2025.12.31.697241

**Authors:** Kiet T. Phong, Siyou Song, Esther Kim, Danny Conrad, Zev Gartner

**Affiliations:** Department of Pharmaceutical Chemistry, UCSF; Department of Surgery, Division of Plastic and Reconstructive Surgery, UCSF

## Abstract

Testosterone is the most studied androgen in hormone replacement therapy in postmenopausal women. Many transgender men also receive long-term testosterone replacement therapy (TRT) and have high serum testosterone levels compared to the low levels seen in cisgender women. Compared to other sex hormones such as estrogen and progesterone, the effects of testosterone on the mammary gland have been relatively understudied and there is little data regarding the long term safety of this treatment. Comparison of mammary glands from transgender men on TRT and cisgender women can reveal the effects of testosterone treatment on mammary gland biology and provide critical information regarding the long-term effects of TRT on patient health and disease outcomes. In this study, we performed single-cell RNA sequencing of breast tissues from a demographics-matched cohort of cisgender women and transgender men on TRT. Surprisingly, participants on TRT had unchanged serum levels of estradiol compared to controls. Among the observed transcriptional differences for participants on TRT were a dramatically reduced expression of genes downstream of estrogen signaling pathways in hormone receptor positive (HR+) luminal epithelial cells, as well as a decreased overall menstrual cycle-related hormone signaling. We confirmed this finding experimentally by showing reduced expression of progesterone receptor a/b, a prominent marker of estrogen signaling, in donors on TRT. Our results support the hypothesis that high levels of testosterone in transgender men on TRT suppress sex hormone signaling in the breast as seen on their impact on HR+ mammary epithelial cells, with implications for TRT as an antagonist of estrogen signaling and protection against breast cancer.

## Introduction

Estrogens and testosterone are the primary sex steroid hormones and have complex and varied impact on the mammary gland where they can either stimulate or inhibit epithelial proliferation depending on the context^1^. Estradiol is produced in the ovaries and play an indispensable role in mammary gland development and physiology as the most potent of the physiologic estrogens during reproductive life^2,3^. Serum estradiol level also rises and falls across the menstrual cycle, and in the mammary gland it activates estrogen receptor alpha (ERα) in hormone receptor-positive luminal epithelial cells to initiate a complex series of paracrine signals that control cell proliferation, differentiation, and tissue morphology ^3^. Increased exposure to ovarian hormones correlates with increased risk of developing breast cancer, for example through an increased number of years in hormone cycling^4^.

Androgens, such as testosterone, also play many important roles in the well-being of women, exerting effects on both reproductive and non-reproductive tissues^5,6^. Although no current FDA-approved testosterone formulation has been approved for use in post-menopausal women, clinical trial suggest that low-dose testosterone can lead to improvements in select patients diagnosed with hypoactive sexual desire disorder. However, the long-term safety of this treatment is unclear, particularly when not used in conjunction with estrogens therapy^5,7^ . Testosterone is the precursor of all estrogens through the activity of aromatase, which is widely expressed in many tissues including breast epithelium and adipose tissues. Testosterone can also signal through the androgen receptor (AR), which is expressed more broadly compared to the ERα^1^. High testosterone levels, together with high aromatase activity, can therefore lead to increased local tissue availability of estrogens and cell proliferation^1^. However, experiments using cancer cell lines and mouse models suggest that androgen treatment can reduce epithelial proliferation and is protective against breast cancer^8–10^. Thus, the role of testosterone in breast cancer is complex, with potentially conflicting or context-dependent effects.

Testosterone is also commonly used in hormone replacement therapy for transgender men. Transgender individuals are people whose gender identity differ from their sex assigned at birth. In recent years, there has been increased recognition and accounting for the number of people who identify in the trans umbrella^11^, with over 1.4 million adults identifying as transgender in the US in 2022^12^. Of that population, about 35.9% identify as transgender men, which refers to individuals who were assigned female at birth but have a male/masculine gender identity. Many transgender men undergo long-term hormone replacement therapy with testosterone to achieve a masculine phenotype, which include changes to body compositions and muscle and fat distributions, development of facial and body hair, as well as cessation of menstruation, which is reported in 86-97% of participants^13^. The serum levels of testosterone in these individuals are much higher than that of cisgender women, and can be in the range of cisgender men^14^ . Since there is evidence supporting a role for testosterone in both promoting and inhibiting pathways associated with ER+ breast cancers, the impact of testosterone replacement therapy (TRT) on the breast in transgender men is unclear^8^. Breast cancer is one of the most common cancers in cisgender women, and thus their screening and care recommendations are made based on extensive studies. On the other hand there is a lack of breast cancer risk data specific to transgender men, making it difficult to provide evidence-based care for this population^15–18^. While many transmen may elect to undergo gender-affirming mastectomy, many may not for a variety of reasons including preference, medical comorbidities, and limited healthcare access. Gender-affirming mastectomy has different goals than prophylactic mastectomy, and may not provide the same level of protections against breast cancer incidence^19^. Therefore, there is an additional need for research to elucidate the effects of androgen treatment on the mammary gland in transgender men relative to breast cancer risks, and specifically on how testosterone modifies the tissue exposure and response to estradiol.

Bentz et al^20^ performed bulk RNA microarray of a cohort of donors before and after TRT, and identified differentially-regulated genes after testosterone treatment. However, the data acquisition method does not account for changes in cell type composition, given that transgender men on testosterone have been reported to show various degrees of lobular atrophy and fibrous stroma^21^. More recently, Raths et al^22^ established an important single-cell resource on the molecular effects of testosterone and described a variety of changes across cell types present in the breast, including changes to gender-specific gene expression, laminin production, and metabolic activities.

Here we build on this previous work to establish a cohort of breast tissues to study the effects of TRT on the mammary gland. Participants in this cohort were selected to have similar parity status, ages, BMIs, and other demographics parameters. We performed single-cell RNA sequencing (scRNA-seq) on the mammary tissue from donors with and without testosterone treatment and performed immunofluorescence staining to validate the results from transcriptomic analysis. Among our major findings, we reveal that the hormone receptor-positive (HR+) luminal epithelial cells in patients on TRT have a reduced amount of luteal phase-related hormone signaling and particularly the estrogen signaling pathways. Protein staining of estrogen and progesterone receptors validated this finding. Overall, our analysis shows that in transgender men, TRT reduces signaling downstream of estrogen in the mammary gland. This finding suggests that the high concentration of testosterone in these donors does not lead to increased exposure to estradiol from local aromatase activities in the mammary gland.

## Methods

### Sample collection

Human breast tissues were collected from reduction mammoplasty or gender-affirming mastectomy surgeries performed at University of California, San Francisco. This study was approved by the Institutional Review Board of UCSF (IRB number 18-25800), and consent and a demographics questionnaire were obtained from all donors prior to collection. After surgical resection, tissues were placed in cold collection medium (1x Antibiotic-antimycotic (Gibco 15240062), 50 U/mL polymyxin B (Sigma P4932), 2% charcoal-dextran stripped FBS (Gemini Bio-Products 100-119) in phenol red-free DMEM/F-12 (Gibco 21041025)) and processed within 2-3 hours. From each donor, approximately 5 mL of blood was drawn and centrifuged at 800 x g for 10 min to collect plasma. Plasma testing was done at Quest diagnostics for total testosterone (MS test 15983), estradiol (ultrasenstitive LC/MS 30289), and progesterone (LC/MS 2839-9).

### Tissue processing

Breast tissue samples were processed into epithelium-enriched tissue microfragments according to LaBarge et al ^23^ before cryopreservation. Briefly, the tissues were minced into a slurry before overnight enzymatic digestion at 37 °C in a spinner flask with digestion medium: phenol red-free DMEM/F12 (Gibco 21041025) containing 200 U/mL collagenase type 3 (Worthington CLS-3), 100 U/mL hyaluronidase (Sigma H3506), 1x Antibiotic-antimycotic (Gibco 15240062), 50 U/mL polymixin B (Sigma P4932), and 10% charcoal-dextran stripped FBS (Gemini Bio-Products 100-119). The digested tissues were centrifuged at 400 x g for 10 min, resuspended in phenol red-free DMEM/F12 medium containing 2% stripped FBS, and passed through serial filtration with 100 µm and 40 µm nylon mesh strainers to separate large tissue microfragments, medium size organoids, and single cells. After size separation, each tissue fraction was cryopreserved in cell freezing medium (RPMI 1640, 40% stripped FBS, 10% DMSO) and maintained in liquid nitrogen cell storage.

### Single-cell RNA sequencing

The tissue microfragments (>100 µm) were thawed and digested in 0.05% trypsin at 37 °C for 2-5 minutes, followed by treatment with 5 U/mL dispase (Stem Cell Technologies 07913) and 1 mg/mL DNase (Stem Cell Technologies 07900) for 2 minutes at room temperature. The suspension was neutralized in PBS + 2% stripped FBS and passed through a 40 µm cell strainer to obtain single-cell suspensions. Cells from each donor were then labeled with donor-specific MULTIseq oligonucleotides (McGinnis 2019^24^ and Murrow 2022^25^) before being pooled. The pooled cells were then sorted on a BD FACSAria III Cell Sorter for viable cells using DAPI as a live/dead indicator. Single-cell gene expression libraries were generated with the Chromium Next GEM Single-cell 3’ Kit (v3.1, 10x Genomics) following standard protocols with some modifications to also generate the MULTIseq barcode libraries. Library concentrations and quality controls were performed on High Sensitivity DNA Bioanalyzer Chips (Agilent Technologies 5067-4626) and Qubit dsDNA HS Assay Kit (Thermo Fisher Q32854). The transcriptome and barcode libraries were pooled and sequenced with a NovaSeq 6000 system on a S4-200 chip (Illumina), targeting 50,000 reads per cell for the transcriptome libraries and 3,000 reads per cell for the barcode libraries.

### Data processing and demultiplexing

The transcriptome fastq files were preprocessed using 10x Genomics Cellranger v5.0.0 and aligned to the human reference genome GRCh38-3.0.0. Cell barcodes were defined as those associated with >630 RNA UMIs and <10% of reads being mitochondrial genes to exclude empty droplets and low-quality cells (Figure S1b-c). Sample demultiplexing was performed on the MULTIseq barcode libraries using the deMULTIplex2 pipeline (Zhu et al 2024^26^) that assigns each cell barcode to a single donor. To validate the deMULTIplex2 classifications, the cellsnp-lite^27^ and Vireo^28^ pipeline were used to assign cells based on donor single nucleotide polymorphisms (SNPs). Analysis of the sequencing data was performed in RStudio (Posit Software, build 561) using Seurat v5.1.0 package^29^. Perturbation analysis was performed with the R package Aurgur^30^ on a down-sampled dataset to minimize error from cell count mismatch between donors (see Figure S4).

### Immunofluorescence

Mammary tissue microfragments were fixed in 4% paraformaldehyde for 2 hours at room temperature and then embedded in 1% agarose blocks in Tris-Acetate EDTA buffer (Sigma-Aldrich 574797). The blocks were placed in a sequential sucrose infiltration gradient up to 30%, then immersed in OCT before transferring into a cryomold and frozen down in an isopentane-dry ice (Sigma 320404) slurry. The frozen blocks were then sectioned at 5-7 µm thickness on a cryostat (Thermo CryoStar NX70). For IF staining, the slides were air-dried at room temperature for 30 minutes before rehydrating in PBS, permeabilized with 0.2% Triton-X, and antigen retrieval was performed in a citrate buffer (10 mM citric acid, 28 mM NaOH in water) in an electric pressure cooker (Instant Pot IPLUX60 V3) on high-pressure steam setting for 3 minutes before cooling down under tap water. The sections were then blocked with 10% normal goat serum in PBS for one hour before overnight incubation with primary ERa antibody at 4 °C. Sections were washed three times with PBS + 0.05% Tween (PBS-T) and tyramide signal amplification (TSA) protocol was applied per manufacturer’s instructions. After TSA staining, the sections go through another round of antigen retrieval to remove the first primary antibody, undergo an additional round of blocking and primary antibody incubation for PRa/b and Cytokeratin 19, followed by TSA staining against the PRab primary antibody. Finally, sections were incubated with DAPI and secondary antibody against Cytokeratin 19 primary antibody for 1 hour before washing three times with PBS-T, and coverslips were mounted with VectaShield Vibrance Antifade mounting medium (Vector Laboratories H-1700). Immunofluorescence imaging was performed on a Zeiss Axio Observer Z1 spinning disk confocal microscope with a Yokagawa spinning disk system.

Antibodies and reagents used are as follow: ERα (1:500; Abcam ab16660), PRa/b (1:100; Cell Signaling Technology 8757S), Cytokeratin 19 (1:200; BioLegend 628502), Alexa Fluor™ 555 Tyramide SuperBoost™ Kit (Invitrogen B40923), Alexa Fluor™ 647 Tyramide SuperBoost™ Kit (Invitrogen B40926), normal goat serum (Gibco 16210064).

### Quantification of hormone receptor expression

Image segmentation was performed in ilastik (v1.3.3)^31^ with the pixel classification pipeline to generate masks for the luminal cell marker KRT19 and the hormone receptors ERα and PRa/b. Then the luminal cell mask is applied to the original image, which subsequently goes into the cell density counting pipeline to estimate the number of luminal cells. Quantification and statistical analyses are performed in Fiji^32^ and RStudio (Posit Software, build 561).

## Results

### Testosterone treatment most strongly affects the transcriptome of hormone receptor-positive luminal cells

We collected breast tissue from 14 donors undergoing either reduction mammoplasty or gender-affirming mastectomy, with 7 donors on active testosterone replacement therapy (TRT) and 7 donors without TRT. All donors are nulliparous and were selected to have matched age and BMI ranges (Table 1). No donors had a known history of breast cancer at the time of surgery. Donors who are on TRT had been in treatment for 18.4 (± 6.78) months on average, and their plasma total testosterone levels range from 234-758 ng/dL. Total plasma testosterone levels of those not on TRT are from 11-42 ng/dL. For comparison, the cis male range is 300-800 ng/dL and the cis premenopausal female range is <50 ng/dL. There is no significant difference in the plasma estradiol level between the donor groups. Most donors not on TRT report regular menstruation (except EK011 and EK035), while almost all donors on TRT report that they don’t have regular menstrual cycles. Most donors on TRT use testosterone cypionate, which is usually a regular intramuscular injection.

**Table 1:** Demographics information and hormone measurements of the participants in this study.

| Donor | Age | BMI | Testosterone treatment | Time on testosterone (months) | Estradiol, plasma (pg/mL) | Progesterone, plasma (ng/mL) | Total testosterone, plasma (ng/dL) |
| --- | --- | --- | --- | --- | --- | --- | --- |
| EK007 | 26 | 31 | No |  | 30 | <0.1 | 13 |
| EK011 | 20 | 26.6 | No |  | 28 | <0.1 | 24 |
| EK013 | 24 | 25.09 | No |  | 106 | 0.3 | 21 |
| EK024 | 19 | 23.83 | No |  | 95 | 7.8 | 11 |
| EK031 | 18 | 29.95 | No |  | 104 | 0.1 | 22 |
| EK034 | 25 | 20.35 | No |  | 171 | 4.4 | 42 |
| EK035 | 24 | 28.84 | No |  | 21 | <0.1 | 31 |
| EK004 | 20 | 23 | Yes | 24 | 12 | <0.1 | 384 |
| EK005 | 27 | 23.8 | Yes | 10 | 24 | <0.1 | 234 |
| EK008 | 19 | 25.8 | Yes | 15 | 67 | <0.1 | 389 |
| EK009 | 21 | 17.4 | Yes | 16 | 25 | <0.1 | 758 |
| EK012 | 31 | 24.85 | Yes | 20 | 34 | <0.1 | 618 |
| EK028 | 24 | 24.96 | Yes | 14 | 38 | <0.1 | 507 |
| EK029 | 20 | 33.2 | Yes | 30 | 87 | <0.1 | 357 |

To investigate the effects of testosterone on the mammary gland, we performed single cell RNA sequencing on breast tissues from the 14 donors using the 10x Chromium platform (Figure 1A). We pooled samples using MULTIseq barcoding^24^ to multiplex samples and reduce batch effects. Input cells were selected for live singlets (DAPI-gate) by FACS without enriching for any tissue compartment to capture the full complexity of the breast tissue (Figure S1A). After filtering for cell quality metrics, demultiplexing with deMULTIplex2^26^ and the SNP-based demultiplexing algorithm Vireo^28^ (Figure S1B-C), the dataset contains 24,544 cells that resolve into 12 distinct clusters (Figure 1B-D) that are annotated based on expressions of known cell type markers (Figure S2). The epithelial clusters are annotated as HR+ luminal cells, secretory luminal cells, and myoepithelial cells. The stromal clusters include fibroblasts, endothelial cells, lymphatic endothelial cells, smooth muscle cells, and vascular smooth muscle cells. Finally, the immune clusters comprise T/NKT cells, B cells, and plasma cells.

**Figure 1:**
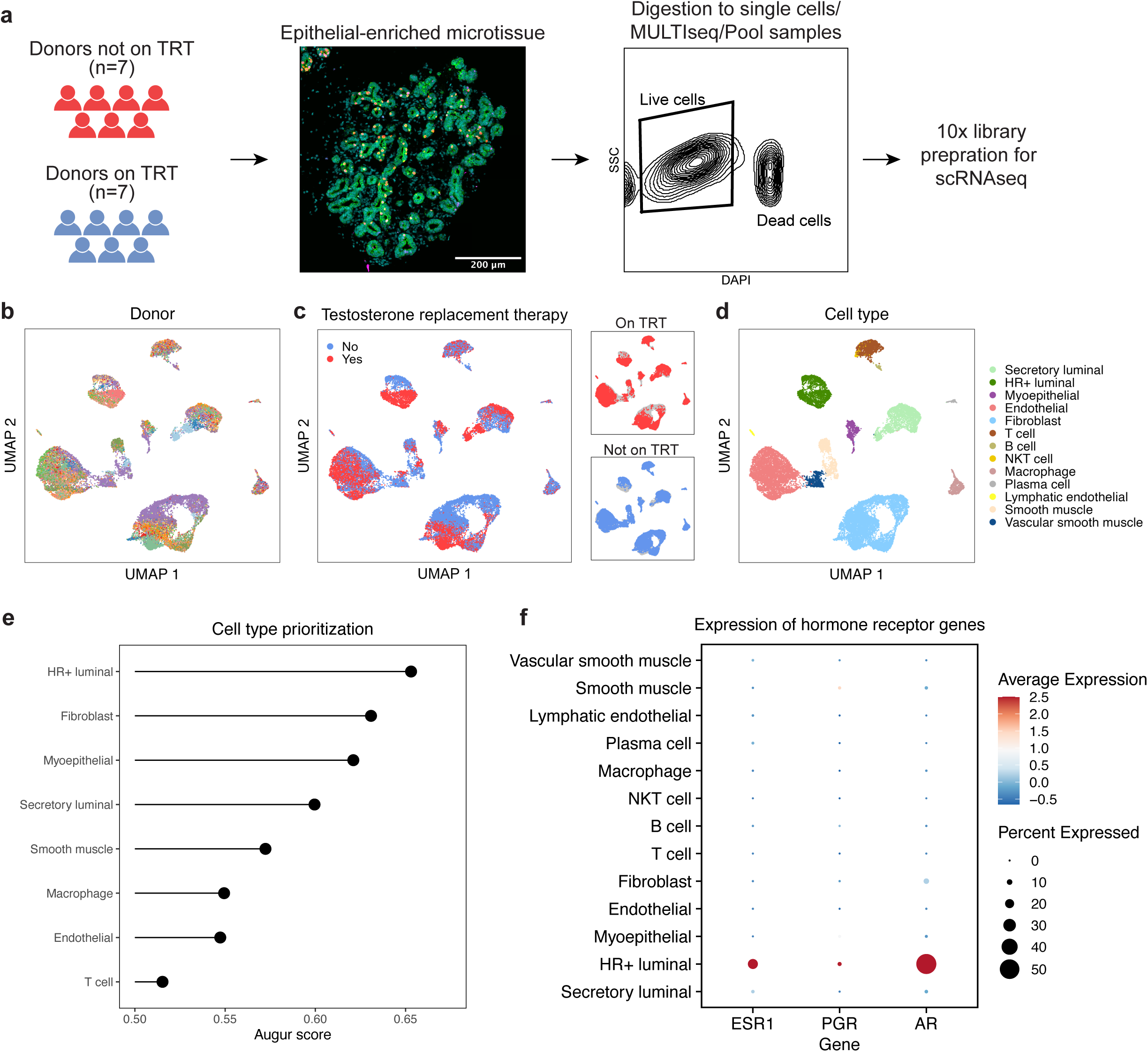
Single-cell RNA sequencing of breast tissue in a cohort comprising healthy donors receiving testosterone replacement therapy (TRT) and controls. a. Tissue processing pipeline from donor sample to 10x library prep. b. UMAP embedding of the demultiplexed scRNAseq dataset colored by donor. c. UMAP embedding of the dataset colored by testosterone replacement therapy status. d. UMAP embedding of the dataset annotated for the major epithelial and stromal cell types in the mammary gland (see Figure S2 for the marker gene panel). e. Cell type prioritization with Augur on a downsampled dataset shows that hormone receptor-positive (HR+) luminal cells have the highest AUC score, indicating that these cells have the largest transcriptomic changes based on testosterone treatment status. f. Dot plot of the gene expression of the hormone receptors estrogen receptor alpha (*ESR1*), progesterone receptor (*PGR*), and androgen receptor (*AR*) among the cell types in the dataset across all the participants, showing that HR+ luminal cells are the cells expressing these receptors on the transcriptome level.

We first investigated whether hormone treatment influences the proportion of cell states within each cell type. We found differential representation of hormone-treated and untreated donors in each cluster (Figure S3). However, the direct comparison of cluster distribution is not reliable because certain cell types and donors are over-represented in terms of cell numbers, which could have resulted from uneven tissue sampling and dissociation. For example, we observed a disproportionate number of fibroblasts from donor EK031 compared to all other donors, and these cells form a large subcluster among the fibroblasts. In order to perform a more balanced quantification of the effects of testosterone treatment on each cell type, we applied the cell type prioritization algorithm Augur^30^ to our data, with TRT as the perturbation. The Augur algorithm uses a random forest classifier to classify cells within each cell type based on the perturbation status and assigns a score based on how well the classifier performs, with a higher score indicating that cell type is more strongly affected by the perturbation. To reduce the influence of cell count imbalances between donors, we down-sampled the number of cells of a given cell type from each donor to the average cell count of that cell type, and then sampled an equal number of cells for each category of testosterone treatment (Figure S4A-B). The Augur analysis revealed that testosterone treatment has the strongest effect on HR+ luminal cells, followed by fibroblasts, myoepithelial cells, and secretory luminal cells (Figure 1E). Additionally, the hormone receptor genes such as estrogen receptor alpha (*ESR1*), progesterone receptor (*PGR*), and androgen receptor (*AR*) are mostly expressed within the HR+ luminal population, and *AR* is present in higher percentage of cells compared to *ESR1* and *PGR* (Figure 1F). This expression pattern suggests that the HR+ luminal cells are the most likely to respond to the presence of estradiol and testosterone compared to other cell types. These results together motivated further investigation into specific transcriptional changes within the HR+ luminal population.

### HR+ luminal cells show decreased signatures of hormone signaling

HR+ luminal cells play a major role in sensing estradiol/progesterone levels and tissue responses through the release of a variety of paracrine signals. Therefore, changes to the hormone microenvironment are expected to have a strong and direct impact on these cells. The UMAP embedding of HR+ cells shows clear separation of populations based on testosterone treatment status (Figure 2A), with decreased expression of many genes related to the estrogen signaling pathway in testosterone treated donors (Figure 2B). In particular, amphiregulin (*AREG)*, an essential mediator of estrogen signaling in mammary gland development^2,3^, is down-regulated in donors on TRT. Wnt Family Member 4 (*WNT4)* and tumor necrosis factor ligand superfamily, member 11 (*TNFSF11*/*RANKL)*, which are major signaling nodes downstream of progesterone and are important paracrine factors to progesterone-induced proliferation and side-branching of the mammary epithelium in vivo, also exhibited decreased expression (log2 fold change are -3.31 and -4.47, respectively). The most significantly down-regulated gene in patients on TRT is parathyroid hormone-like hormone (*PTHLH)* (log2 fold change of -6.81 and adjusted p-value of 2.12e-280), which is a key factor in mammary gland development^33^ and is expressed downstream of epidermal growth factor receptor (EGFR), where it is activated by extracellular signal-regulated kinases (ERK) and p38 signaling^34^. Gene set enrichment analysis on the list of differentially expressed genes in HR+ luminal cells shows a negative enrichment score in cells from donors on TRT for pathways related to cell chemotaxis, humoral immune response, G protein-coupled receptor signaling pathway, and cell motility (Figure 2C).

**Figure 2:**
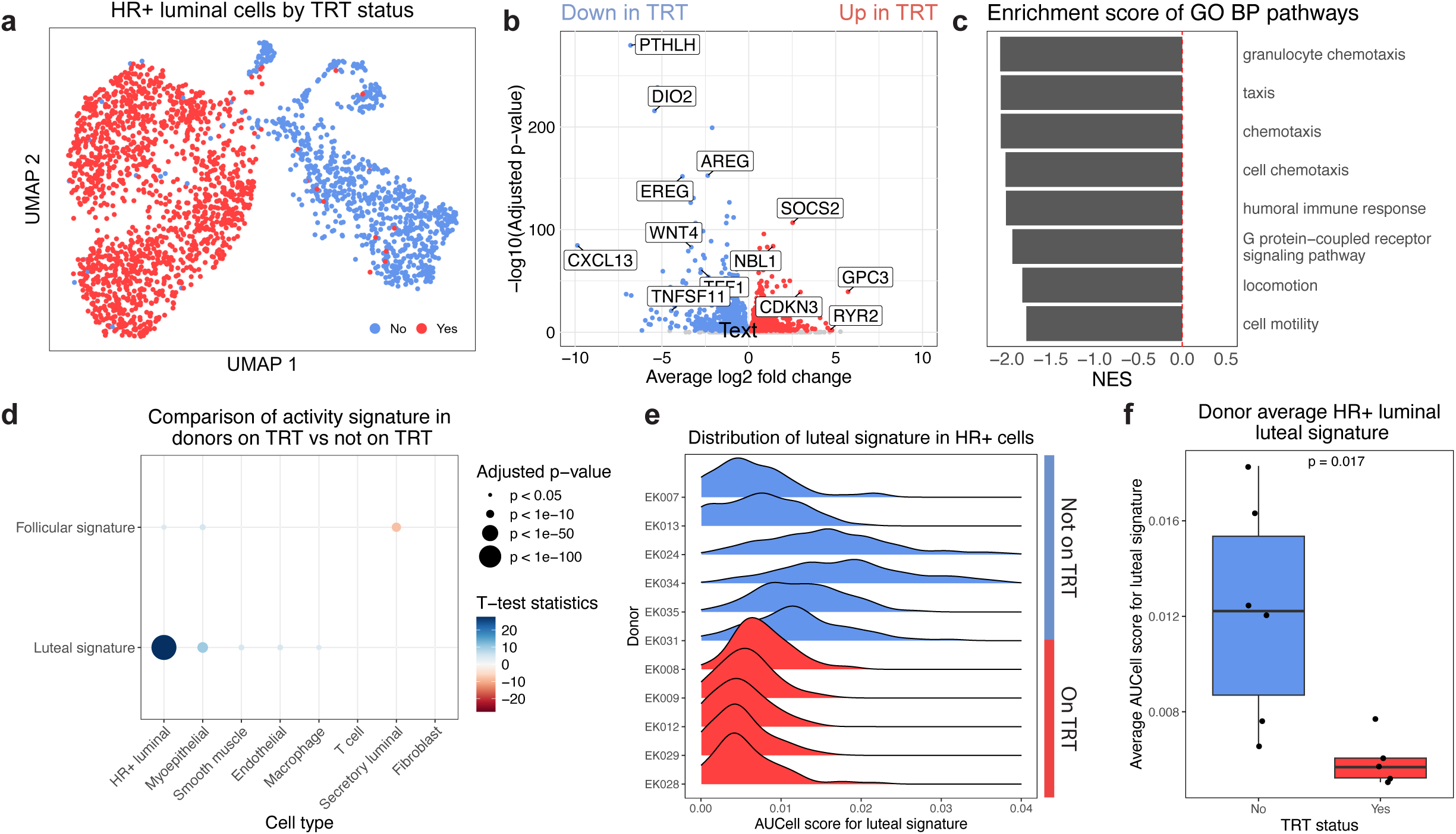
HR+ luminal cells in transgender men receiving TRT have decreased expression of hormone response genes a. UMAP embedding of the HR+ luminal epithelial population colored by TRT status. b. Volcano plot illustrating differential gene expression in HR+ luminal cells based on TRT status and highlighting genes downstream of estrogen signaling. c. Significant GO Biological Process terms and their normalized enrichment scores (NES) based on the fold-changes of the differentially expressed genes between TRT statuses. These terms have negative NES indicating that they are enriched in donors not on TRT and down-regulated in donors on TRT. d. Comparison results of AUCell scores for each cell type and gene set between TRT statuses. AUCell scores for luteal phase activity shows the most significant difference among HR+ luminal cells. P-values and test statistics are produced by student t-test, with Bonferroni correction for multiple comparisons. e. Distribution of the luteal phase activity score in the HR+ luminal cells from each donor f. The average luteal phase activity score in HR+ luminal cells for each donor (as in ‘e’) showing decreased overall scores for donors on TRT.

We then applied AUCell^35^ to calculate the activity score for hormone response signature genes in our dataset. These gene sets are derived from a bulk sequencing dataset of breast biopsies from women at different phases of their menstrual cycle^36^. Given that menstrual cycle-related changes in the mammary gland are driven by the activity of estrogens, progesterones, and their downstream paracrine signaling between various cell types in the tissue, this metric is used as a proxy for quantifying the diverse tissue-level response to these hormones. The advantage of this method is being able to use an empirical metric to quantify the varied downstream effects of these signaling pathways in addition to the direct activation of either estrogen or progesterone pathways that are specific to the mammary gland. When comparing the AUCell score for luteal hormone response activity based on TRT status, HR+ luminal cells show the most significant score difference among all the cell types (Figure 2D). While these gene sets are derived from bulk sequencing of tissues containing multiple cell types, the AUCell score for each gene set is calculated on a single-cell basis. Therefore, given that the HR+ luminal cells only make up a relatively small proportion of the breast, this result highlights that they are the major source of gene expressions that contribute to hormone response activities in the breast.

The distribution of hormone response activity score for control patients shows a wide range of values. Given that hormone levels rise and fall across the menstrual cycle, and scheduling of surgery is independent of menstrual cycle stage, this observation is consistent with the notion that each donor is at a different phase of their menstrual cycle (Figure 2E-F). For example, one would expect that donors who are in the luteal phase of their menstrual cycle will have a higher score for luteal phase gene activity while those in the follicular phase will have a lower score. The donors with the highest plasma progesterone levels and high estradiol levels, EK024 and EK034, also show the highest AUCell score for the luteal signaling pathway, which is consistent with the high hormone environment interpretation of this score. HR+ luminal cells from donors on TRT, in contrast, have uniformly low luteal phase activity scores, which suggest that these cells don’t experience the high hormone signaling environment of the luteal phase. Plasma measurements of these donors also show consistently very low levels of progesterones. Combined, our analysis suggests that the mammary glands from the donors on TRT do not experience luteal phase signaling activity, which is driven by increased serum levels of estrogens and progesterones. This result is consistent with our finding of reduced expressions of signaling factors downstream of estrogens and progesterones activity.

The AUCell method also shows that HR+ luminal cells in donors on TRT has a significantly increased score for the Hallmark androgen response pathway (Figure S5), which is consistent with the donor groups. However, this is a very modest increase, as the pathway was not defined specifically for the mammary gland, and there could be tissue-specific responses to androgen.

Our findings are mostly agree with the results of an independent study by Raths et al, where they performed single nucleus RNAseq and single nucleus ATACseq of frozen breast tissues from cisgender women and transgender men^37^. In their study, the HR+ luminal cells in transgender men show no change in *ESR1* expression, but there is a reduction in its downstream signaling pathways, including *AREG* and epiregulin (*EREG),* as well as parathyroid hormone-like hormone (*PTHLH*, Figure S7). Furthermore, progesterone receptor transcript expression is downregulated in transgender men in their study. Additionally, they also observed an increase in expression of genes downstream of AR like ryanodine receptor 2 (*RYR2)*, which we also observed in this study (Figure 2B). Overall, our results are generally consistent with Raths et al. on the impact of TRT on HR+ luminal cells as well as similar trends in other cell types. Integration of our results with the Raths et al single nucleus RNAseq data shows generally similar distributions for the cell types (Figure S8). However, some discrepancies in findings may reflect differences in the underlying biology or the low power of these two studies. (Table 2). In our cohort, we did not observe any significant change in the expression of various ECM-related genes and pathways in either HR+ luminal cells or other cell types (Figure S9).

### Mammary glands from TRT patients show decreased expression of progesterone receptors a/b

We performed immunofluorescence staining on the mammary tissue of an expanded cohort of donors (11 not on TRT, and 8 on TRT) to confirm the key findings from transcriptomic analyses. Staining was performed on digested microtissues to increase the amount of epithelial tissues for quantification. Among the donors on testosterone, we observed a significant decrease in the staining for PRa/b, and a non-significant increase in ERα staining (Figure 3A). Quantification of the number of PRa/b-positive luminal cells relative to the total number of luminal cells shows that there is a significant decrease in the expression of the hormone receptor in donors on testosterone (Figure 3B). PRa/b is a target for upregulation by the estrogen receptor in the mammary gland^38^, therefore decreased expression of this hormone receptor is consistent with our observation of decreased estrogen signaling pathway genes in donors on testosterone.

**Figure 3:**
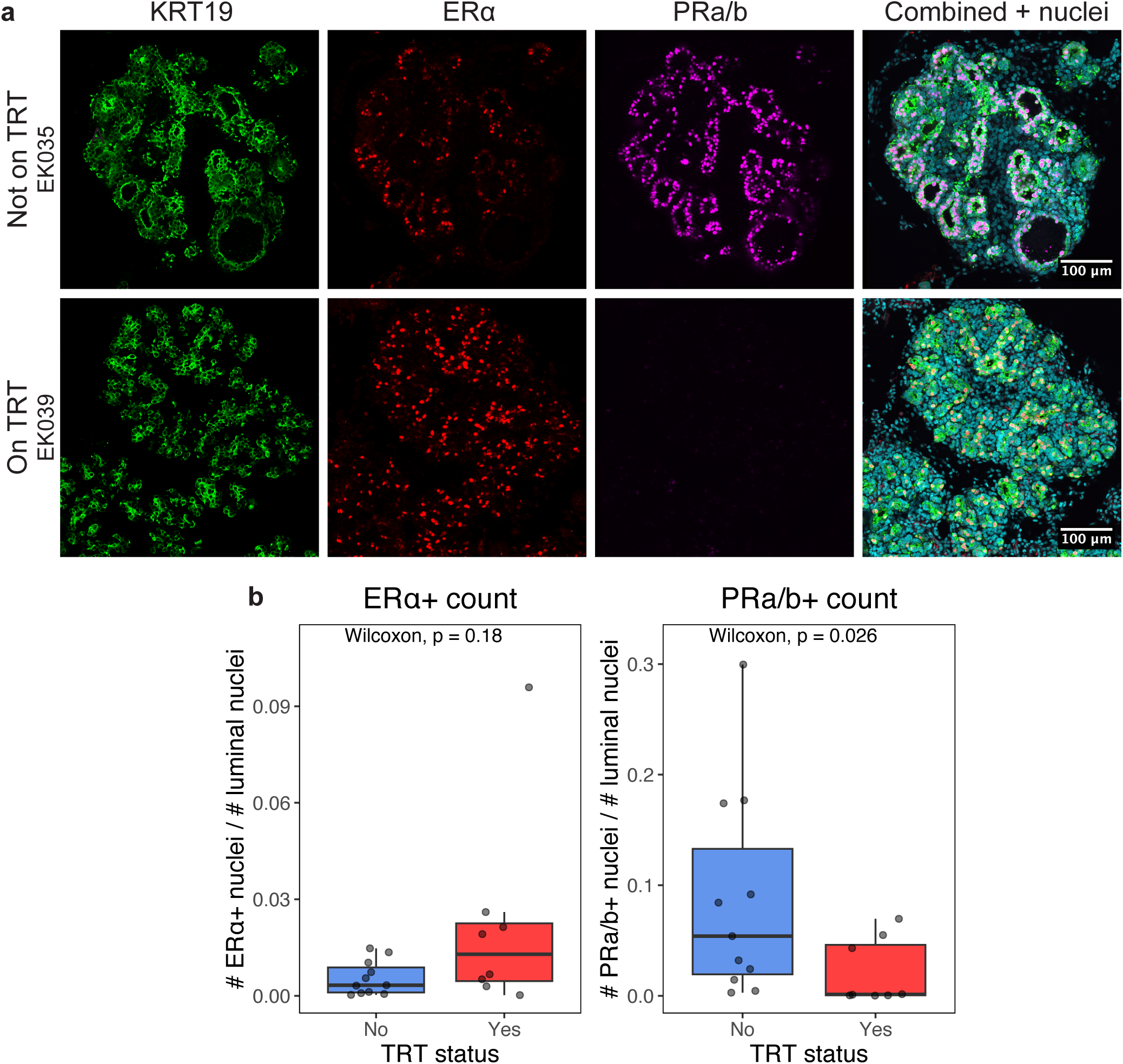
Testosterone treatment reduces PRa/b protein expression in luminal epithelial cells. a. IF staining of partially digested epithelial microtissues for donors with and without TRT. KRT19 (green) is a marker for luminal epithelial cells, and ERα (red) and PRa/b (magenta) are hormone receptors for estrogen and progesterone, respectively. b. Quantification of hormone receptor staining shows minor difference in ERα expression, and a significant decrease of PRa/b expression in donors on TRT.

## Discussion

Androgen exposure and androgen receptor expression are known factors influencing breast cancer prognosis. However, there is conflicting evidence on the influence of androgen on breast cancer progression, likely due to the complexity of interactions between the androgen and estrogen signaling pathways and the cellular context. Testosterone replacement therapy has been used in post-menopausal women to treat hypoactive sexual desire disorder, although no FDA-approved formulation exists due to a lack of long-term safety data. In these patients, the dosage is tuned such that the testosterone level should approximate the pre-menopausal female physiological levels. Additionally, many transgender men undergo testosterone replacement therapy to achieve serum testosterone levels similar to cisgender men, and thus their tissues are exposed to a much higher levels of androgen compared to cisgender women. There is concern regarding the long-term effects of such high testosterone treatment. Current guidelines for breast cancer screening and care for these patients are based on epidemiological data from cisgender women, and there is a lack of data specific to this population. In this study, we have established an independent cohort of donors comprising cisgender women and transgender men to study the impact of testosterone on their breast tissue. These results extend our understanding of the impact of testosterone replacement therapy on the healthy mammary gland, and potentially can provide supporting evidence for the safety of long-term low-dose testosterone treatment in post-menopausal women.

The major finding in our analysis is that there is a decrease of signaling downstream of estrogen in the hormone receptor-positive luminal epithelial cells in the donors on testosterone treatment, as shown by the decreased expression of genes downstream of estrogen receptor such as *AREG*/*EREG*, *PTHLH*, *WNT4*/*RANKL*, and others. AREG is an EGFR ligand that is up-regulated by estrogen, and plays a role in mammary ductal elongation and in driving the proliferation of the mammary stroma^2^. Further downstream, PTHLH is a peptide hormone that has been shown to be is regulated by the EGFR pathway, and blockage of EGFR ligands such as AREG leads to reduced PTHLH expression^33,34^. PTHLH is also involved in the signaling from epithelium to the stroma, particularly in specifying the mammary mesenchyme during development but is also implicated in breast cancer development and bone metastasis^33,34^. Bone metastasis is one of the most prevalent sites of breast cancer metastasis regardless of tumor subtype, where up to 85% of metastatic patients show bone metastasis, and particularly hormone receptor-positive tumors may show higher levels of bone metastasis compared to hormone receptor-negative or triple negative cancer^39^. The impact of PTHLH on prognosis of metastasis disease is complex, where it may be protective against tumorigenesis and cancer growth in the early phase, while promoting metastatic bone disease and reducing median survival rates for late stage cancer^40,41^. WNT4 and TNFSF11 (RANKL) are additional secreted factors that are important mediators of progesterone signaling in the mammary gland and are pivotal in the control of stem cell function and mammary gland development postnatally^42,43^. During normal progesterone signaling in the breast, WNT4 is secreted by the HR+ luminal cells in response to progesterone receptor activity, and activates canonical WNT signaling in the myoepithelial cells, which results in additional changes that feed back to the luminal compartment^43^.

These changes are consistent with a reduction in the activity of the estrogen signaling pathway and its downstream autocrine and paracrine mediators in the mammary gland of transgender men on TRT. Expression of the progesterone pathway is also decreased potentially due to the downregulation of the progesterone receptor because of the reduced estrogen signaling. The luteal phase hormone response signature analysis provides additional support for this model, as this signature score quantifies both the expression of the estrogen and progesterone pathways, and their downstream paracrine interactions with other cell types in the mammary gland. A reduced score in the donors on TRT indicates there is an overall reduced amount of hormone signaling in the HR+ luminal cells, since the luteal phase is characterized by increased circulating levels of estrogens and progesterones. Physiologically, progesterone receptor is up-regulated by estrogen signaling, which primes the mammary gland to respond to an increased level of circulating progesterone during the luteal phase. If there is suppression of the estrogen signaling pathways, then the corresponding decrease of progesterone receptor expression would lead to a decreased activation of the progesterone signaling pathway. Additionally, most of the donors on TRT in our cohort report that they don’t have regular menstrual cycles, and their plasma levels of progesterones are in the range of follicular phase levels. These observations suggest that these donors are anovulatory. Therefore, the systemic impact of testosterone on the endocrine system is a potential mechanism driving the reduced luteal signature in HR+ luminal cells, which no longer receives the high hormone signaling from the luteal phase of the menstrual cycle. Given that these cells are the major source of paracrine signals driving the tissue responses to changing hormone levels during puberty, menstruation, and pregnancy, alterations in the state of the HR+ luminal cells may lead to dramatic changes in both the hormonal microenvironment and the other cell types in the breast that respond to these paracrine signals.

Therefore, the impact of testosterone treatment on the HR+ luminal cells can lead to changes in other cell types that are independent of its activation of the androgen signaling pathways.

Compared with an independent study by Raths et al^37^, our findings in HR+ luminal cells show similar impacts of TRT on gene expression, as well as many changes in other cell types (Table 2). In particular, they also showed an impact of testosterone treatment on *PGR*, *AREG*, and other downstream pathways of estrogen signaling. While each of these two independent studies examined relatively small cohorts, the fact that they both arrived at similar conclusions regarding the major impact of TRT on hormone signaling provides strong support for these findings. Particularly, both our analyses agree on the findings of decreased estrogen signaling in the donors on TRT. Testosterone has been reported to reduce the secretion of ovarian hormones^44^, including estrogens. Thus, the molecular changes we observe could have resulted from reduced circulating estrogen levels or repression of estrogen signaling pathway by testosterone activation of androgen signaling pathways. In our donor cohort, there is no significant difference in plasma levels of estradiol between the TRT groups, which strongly suggests that the reduced estrogen signaling is due to the suppressive effects of testosterone on estrogen receptor-mediated signaling. In vitro stimulation studies of mammary organoids should allow for the isolation of these factors and uncover the specific effects of testosterone on the mammary gland and further validate these findings. However, conditions for in vitro stimulations should be carefully considered, as the effects of androgens on the mammary gland are complex and context-dependent.

One important aspect of testosterone signaling in the mammary gland is that it can be converted into 17β-estradiol by aromatase, which is expressed in mammary and adipose tissues. Particularly after menopause when the ovarian hormone production has decreased, aromatase is thought to be a significant source of estrogens through converting available androgens into estrogens and exerting influence on the local hormonal microenvironment^1,45^. Dysregulation of aromatase has been reported in a variety of breast cancers, and aromatase inhibitor has been evaluated in many clinical trials for treating hormone receptor-positive breast cancer, or as adjuvant to improve survival after tamoxifen treatment^46^ . One major concern about elevated testosterone levels in transgender men on TRT is that increased testosterone levels would lead to local aromatization of testosterone into estradiol, thus exposing their mammary glands to high levels of estrogen signaling. Our data does not support this concern in the context of high testosterone levels, as it points to a reduction of estrogen signaling in the mammary glands of individuals exposed to high levels of testosterone. This could potentially result from aromatase activity being suppressed in these tissues, or that the unusually high abundance of testosterone in donors on TRT counteracts the local production of estradiol by aromatase in the tissue. While there is no evidence for the suppression of aromatase activity by AR signaling, future studies could conclusively refute this possible mechanism by measuring aromatase activity in standardized units of fresh mammary tissues, as well as *in vitro* tests after hormone stimulation. These signaling pathways may also become dysregulated in the context of breast cancer. There have been reports of high AR/ER ratios in the context of high testosterone levels leading to poorer outcomes in pre-menopausal women with breast cancers^47^. These results highlight the complex interplay among these pathways and the need to understand the mechanisms of androgen receptor activity in breast cancer cases in order to provide effective treatment^48^.

ERα and PR are the canonical receptors mediating the activity of estrogens and progesterones in the mammary gland. Our investigations show that there is a non-significant trend of increased ERα expression in donors on TRT in our expanded cohort, and a significant decrease in PR staining. The trend of increase in ERα expression could indicate a complex role of AR activity in modulating the expression of this nuclear receptor. There is conflicting evidence on whether deleting AR in a mouse model will stimulate or inhibit mammary gland development and downstream estrogen signaling^49,50^, potentially due to different activities of the AR domains. On the other hand, PR is commonly used as a biomarker for the activity of estrogen signaling. The decrease in PR expression therefore is consistent with the anti-estrogenic effects of androgen receptor stimulation^9,10^ and demonstrates that there is a reduced estrogen signaling in the mammary gland of donors on TRT.

A limitation of our study is the small size of the donor cohort. Potentially increasing the number of donors in each group would allow a more detailed investigation of the effects of TRT, increasing the statistical power of some comparisons that currently shows a non-significant trend, as well as increasing confidence in the luteal signature analysis. However, integrating our data with the data previously published by Raths et al (Figure S7) shows similar patterns of cell clustering and gene expressions, thus increasing the confidence in our findings. Another drawback is that single-cell sequencing data only captures a static picture of gene expressions, and future stimulation studies in either an organoid or mouse model is needed to determine the kinetics and mechanistic effects of testosterone in the mammary gland.

In conclusion, we established a single-cell dataset of breast tissues from a matched cohort of cisgender women and transgender men. We observed that testosterone treatment produced the biggest effect on the transcriptome of HR+ luminal cells, leading to a decrease in expression of genes in the estrogen signaling pathways and other downstream processes. There is no activation of the estrogen pathway due to local activity of aromatase in the context of high testosterone levels in TRT. These molecular changes, together with systemic changes in hormone levels, result in a decrease in the luteal phase gene activity in these cells in transgender men donors. Increased exposure to hormone cycling through the menstrual cycle has been shown to increase the overall risk of breast cancer^4^, and thus the reduction in hormone signaling could indicate a systemic protective effect of testosterone in this population^8^ and a potential therapeutic target for certain hormone receptor-positive breast cancer subtypes^9,51^. Future studies should validate this finding in larger independent cohorts, dissect the effects of the activation of androgen pathways, and investigate how it could potentially impact carcinogenesis in the mammary gland. Together, these results could create meaningful impact on the clinical care and management for transgender men on long-term TRT, as well as the use of TRT in cisgender women for treating menopause-related symptoms.

## Data Availability Statement

Single-cell RNA-seq data has been deposited at the Gene Expression Omnibus (GSE283161) and will be available upon manuscript acceptance/publication.

## Author Contributions

K.P., Z.J.G, E.K. conceptualized and designed the experiments. E.K. performed the surgeries, and K.P. and S.S. processed the specimens for tissue banking. K.P. and S.S. performed the sequencing experiment. D.C. processed the sequencing data and K.P. performed the data analysis. K.P. performed the immunostaining and analysis. K.P. and

Z.J.G. wrote the manuscript. All other authors reviewed the manuscript and provided feedback. We also thank Vasudha Srivastava, PhD, and Honesty Kim, PhD for reviewing and providing feedback on the manuscript.

## Acknowledgements

This work was funded by the NCI PS-ON grant (U01CA244109) and Department of Defense Era of Hope Expansion Award (W81XWH-13-1-0221) to Z.J.G., the UCSF Center for Cellular Construction (DBI-1548297), and the Barbara and Gerson Bakar Foundation. Z.J.G. is a Chan Zuckerberg BioHub Investigator. FACS was performed on BD Aria 3, supported in part by HDFCCC Laboratory for Cell Analysis Shared Resource Facility through a grant from NIH (P30CA082103). Sequencing was performed at the UCSF CAT, supported by UCSF PBBR, RRP IMIA, and NIH 1S10OD028511-01 grants.

## Supplementary Information

### Supplementary figures

**Supplementary Fig. 1.**
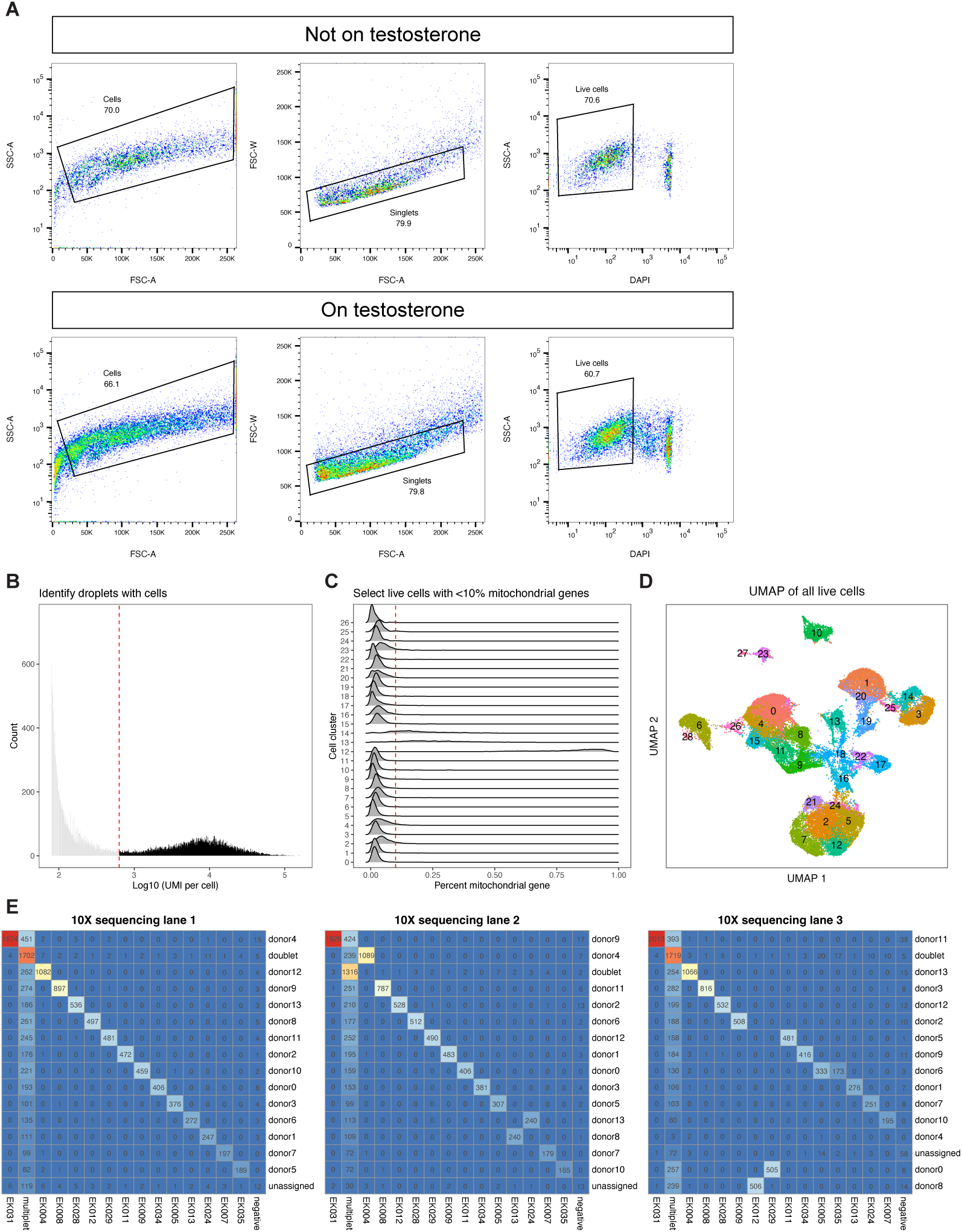
FACS sorting scheme and quality control of single-cell data a. FACS sorting scheme to sort for live cells from each MULTIseq-labeled sample pool. b. A threshold of at least 630 UMI per cell is applied to remove empty droplets. c. Cells with at most 10% of reads coming from mitochondrial genes are kept as healthy cells, while those with higher percentages are removed as unhealthy/dying cells. d. UMAP embedding showing all live cells, used as input for MULTIseq barcode demultiplexing and Vireo donor classification. e. Heatmap comparison of deMULTIplex2 output and Vireo output, showing agreement of most donor classifications and additional classified cells based on Vireo results.

**Supplementary Fig. 2.**
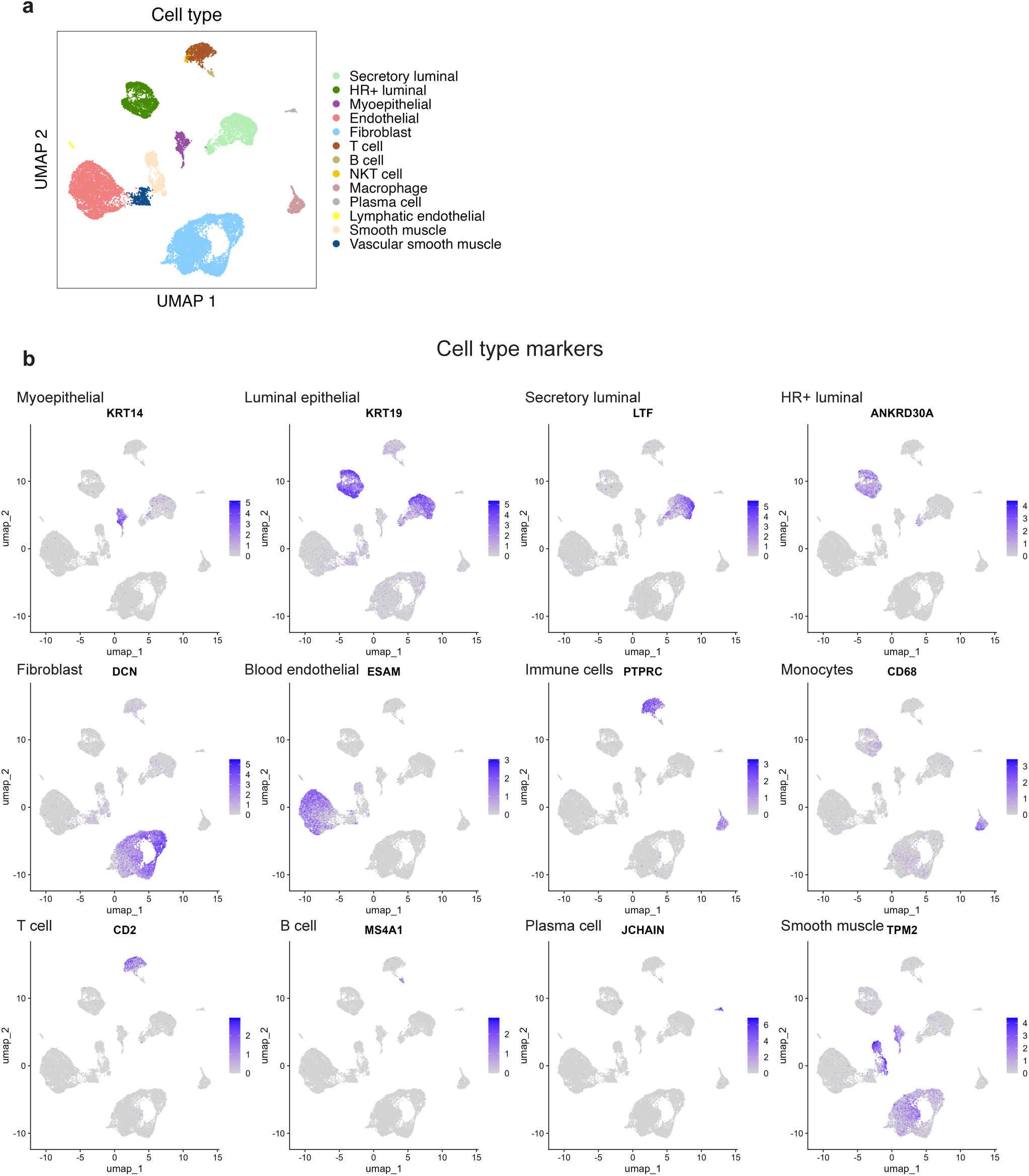
Expression of gene markers used to annotate cell types a. UMAP embedding of demultiplexed cells with the cell type classifications. b. Cell type markers used to classify each cell cluster.

**Supplementary Fig. 3.**
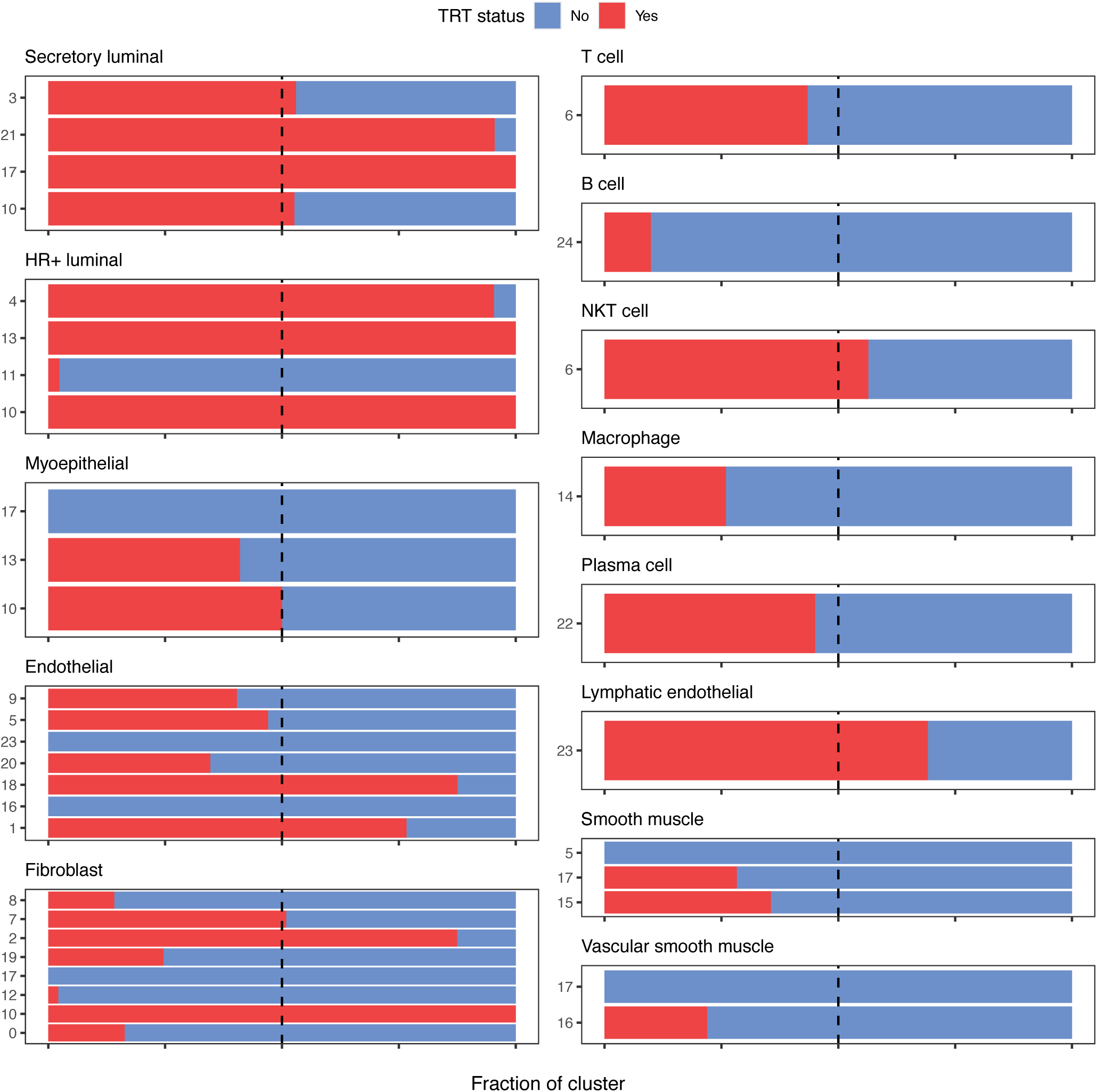
Relative proportion of cells based on TRT status for each cluster and cell type, showing that some clusters are specific to cells from donors on testosterone and vice versa.

**Supplementary Fig. 4.**
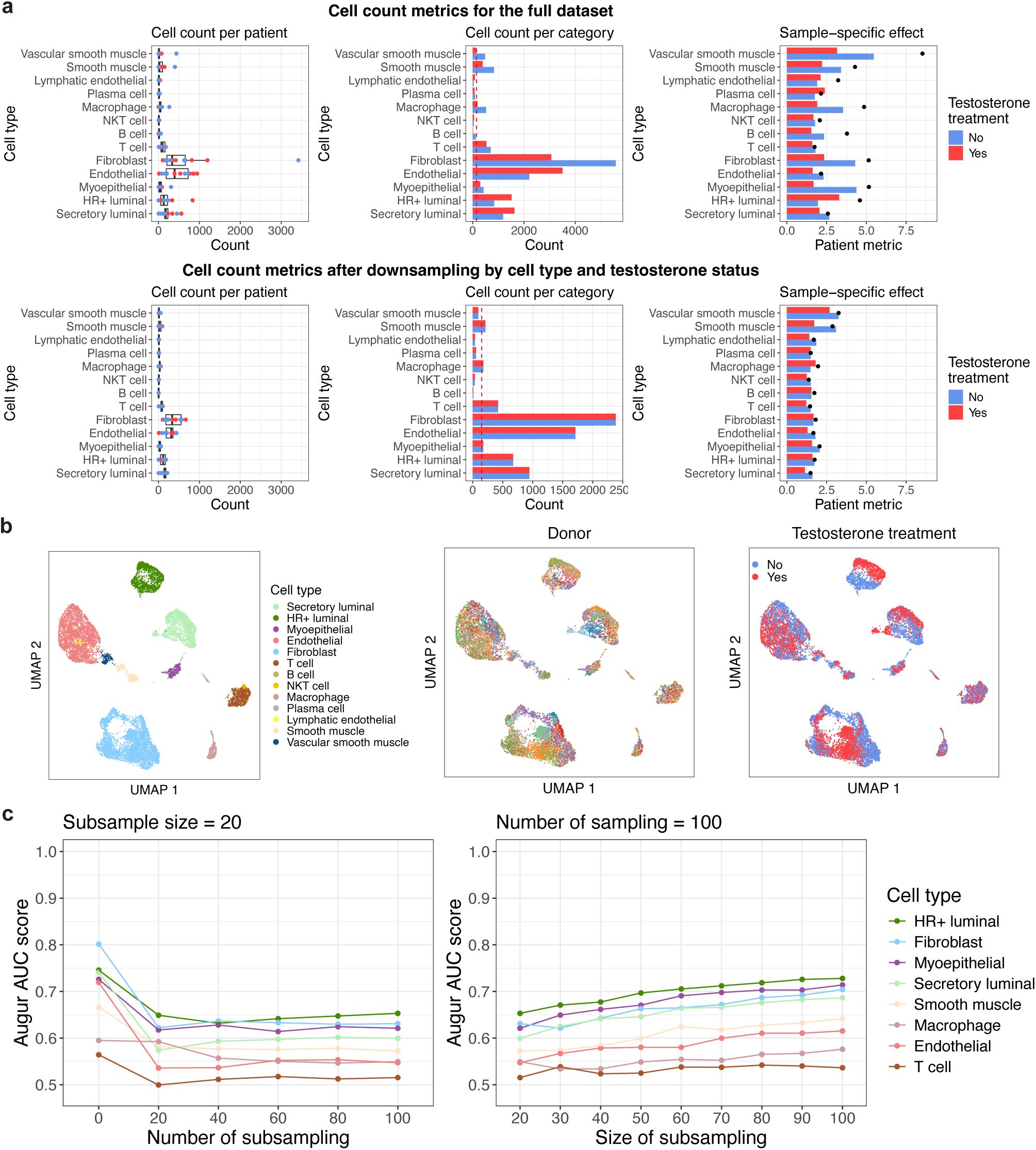
Summary of down sampling strategy for the scRNA-seq dataset to balance cell type number prior to analysis with Augur a. Cell count distributions and patient-specific metrics (patient max cell count divided by mean cell count for each cell type) before and after down sampling. The process of down sampling reduces the influence of the overabundance of cells from a single patient on the Augur score. b. UMAP embedding of the dataset after down sampling showed similar qualitative features as the full dataset. c. Parameter sweep for the Augur algorithm showing stability of the cell type ranking across different values.

**Supplementary Fig. 5.**
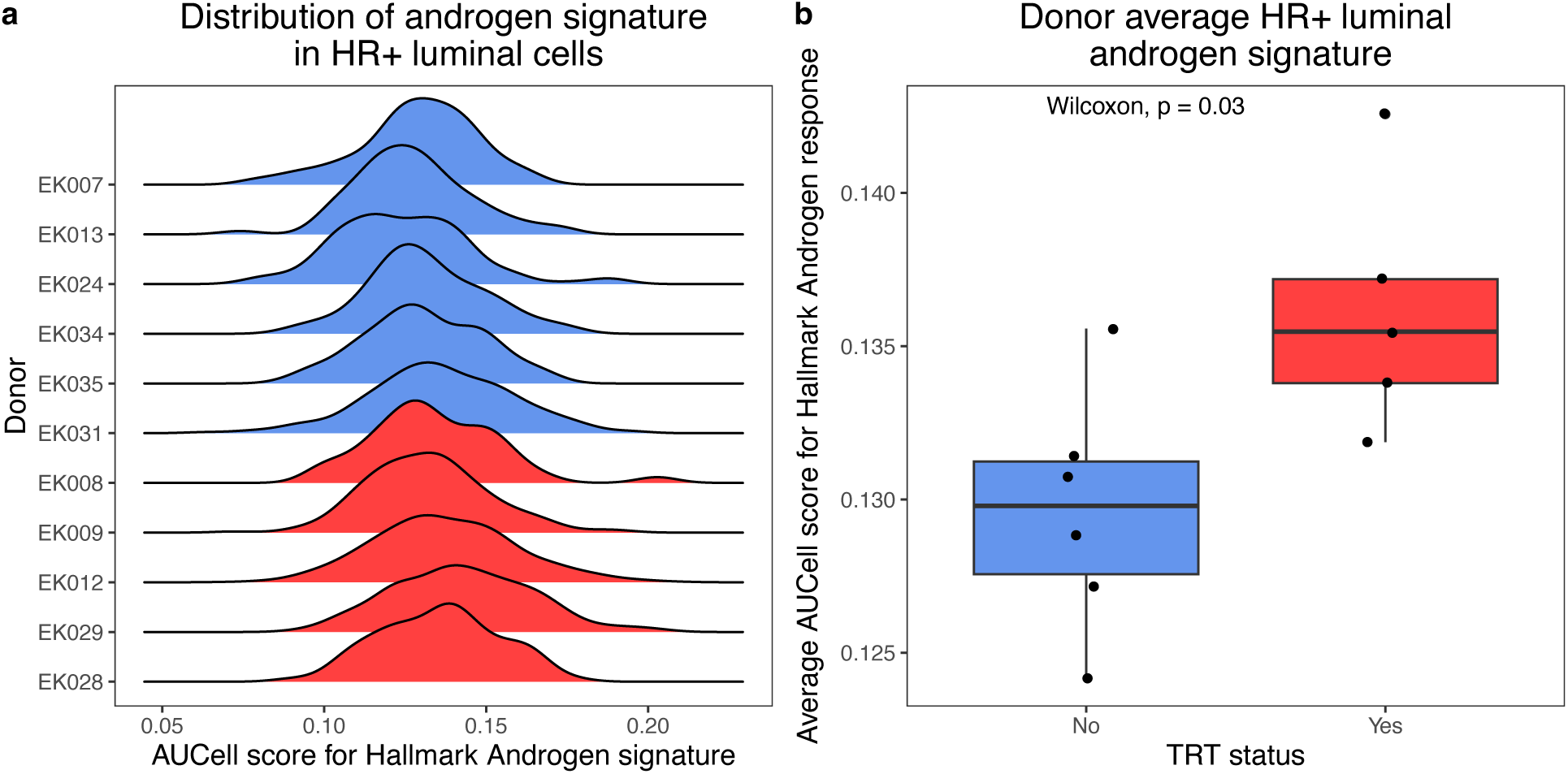
Hallmark androgen response signature in HR+ luminal cells. a. Distribution of AUCell scores for the Hallmark androgen response pathway in HR+ luminal cells. b. Donor average AUCell score in HR+ luminal cells for the androgen signature shows a small but significantly different androgen response in donors on TRT.

**Supplementary Fig. 6.**
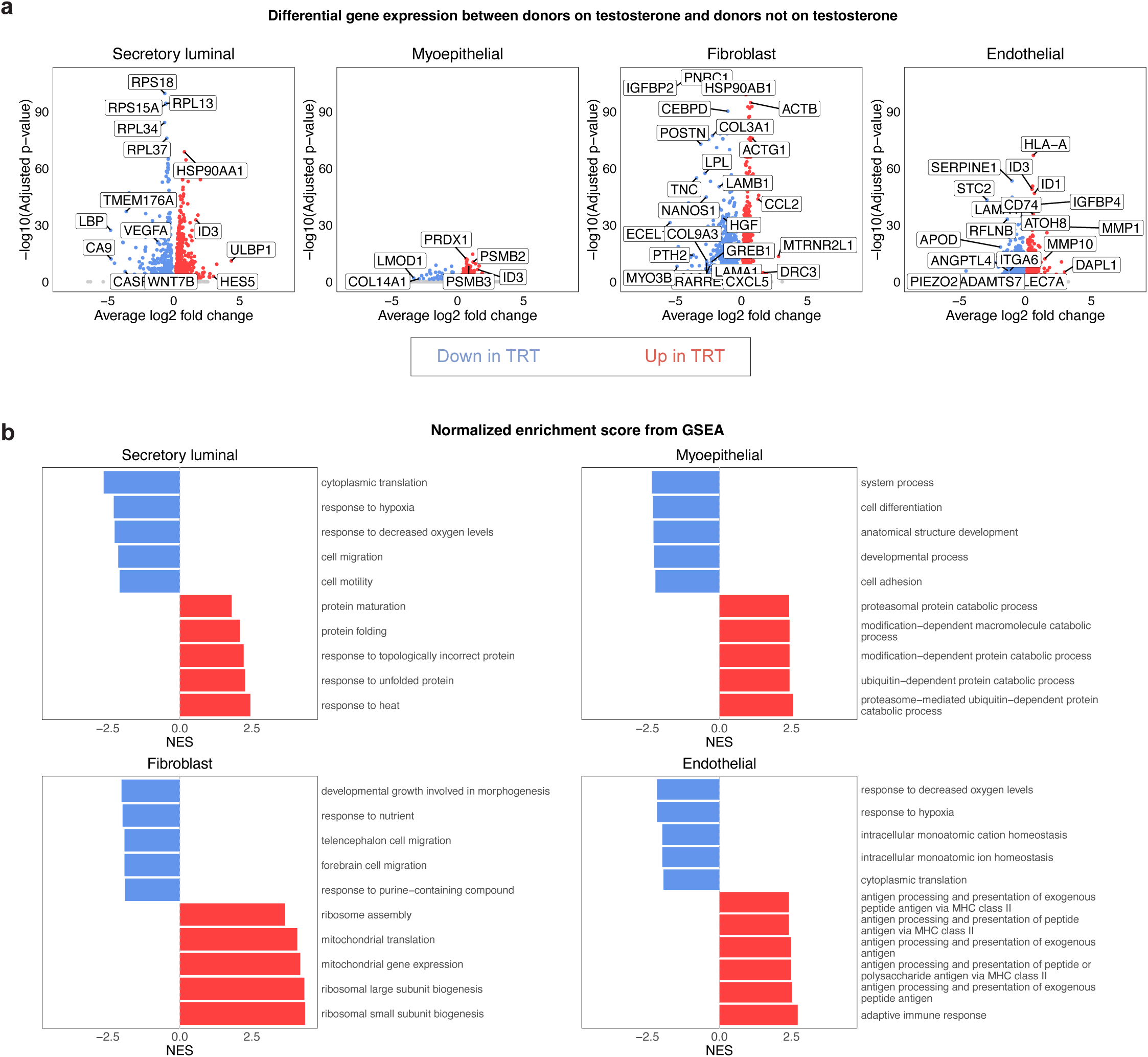
Distinct transcriptional response to TRT in the major cell types of the mammary gland. a. Differentially expressed genes based on TRT status in each major epithelial/stromal cell type. b. Normalized Enrichment Score of the top 10 GOBP pathways in the GSEA results based on the differentially expressed genes. A positive NES means increased enrichment in the donors on TRT, and negative score means increased enrichment in donors not on TRT.

**Supplementary Fig. 7.**
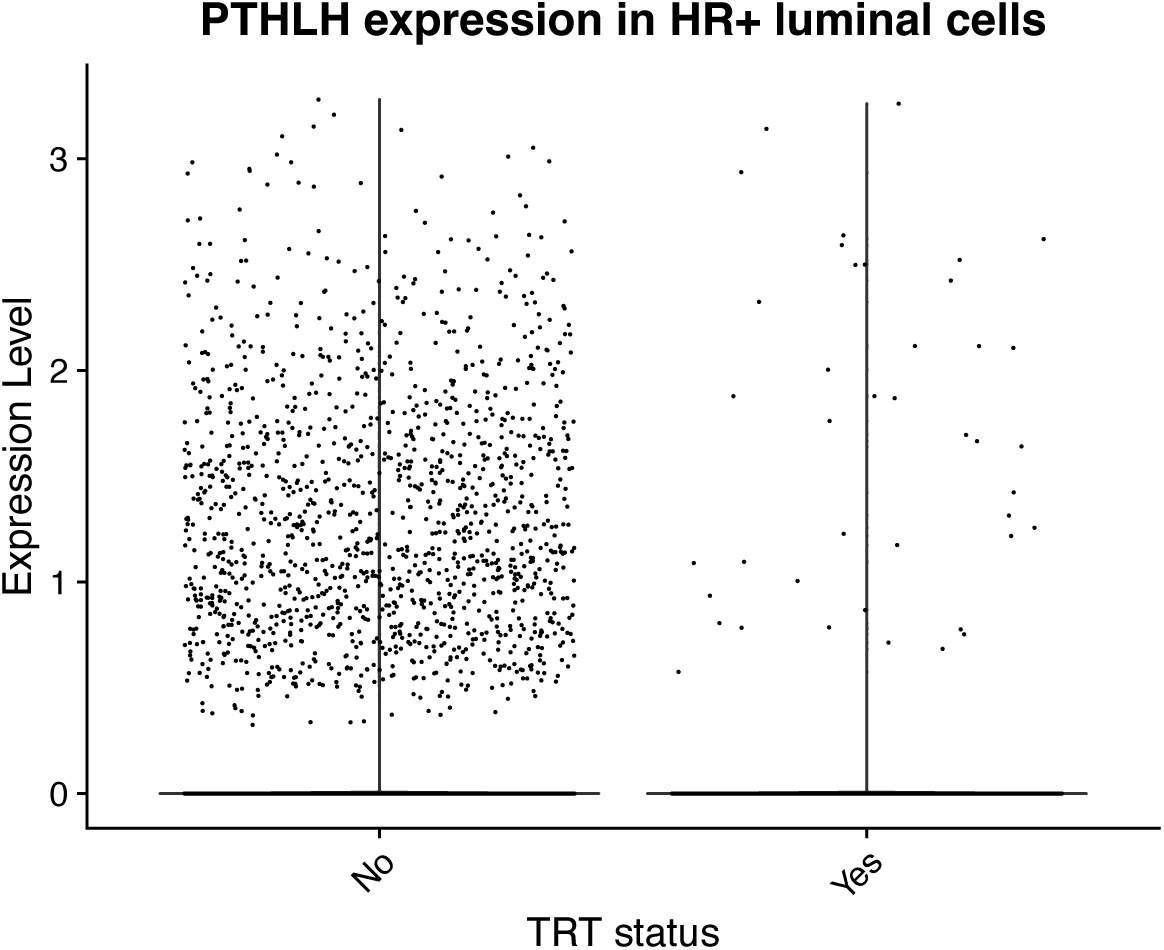
Decreased expression of PTHLH in HR+ luminal cells of participants on TRT compared to those not on TRT in the Raths et al dataset, showing agreement in the molecular finding of hormone-related signaling with the current dataset.

**Supplementary Fig. 8.**
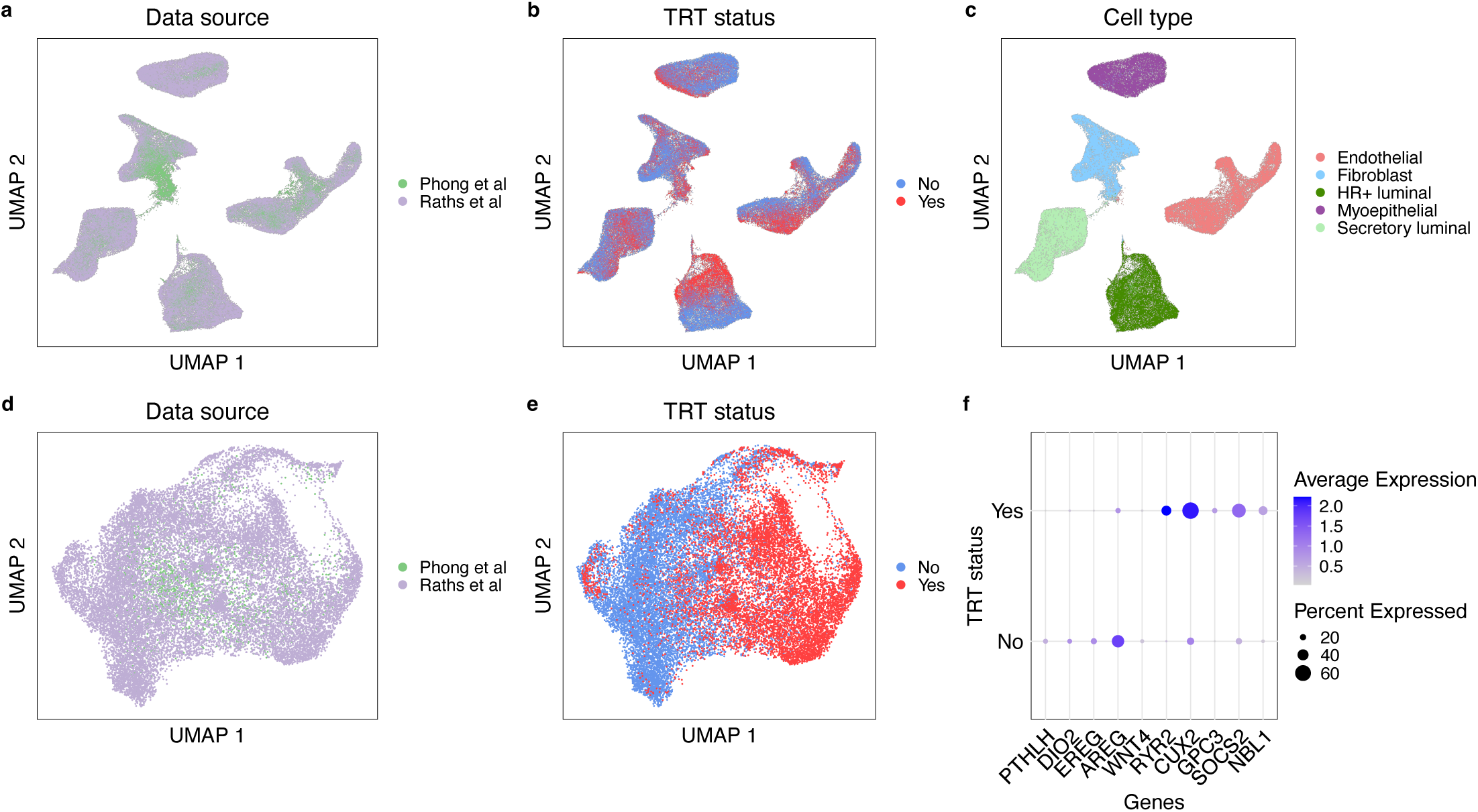
Integration of the single-nucleus RNA seq data from Raths et al and the single-cell RNAseq data from this study. We restricted the data to the three mammary epithelial subtypes, fibroblasts, and endothelial cells, then performed CCA integration to correct for batch effects and technological differences. a. UMAP embedding of the CCA integrated dataset by data source, showing good mixing of most clusters except for the fibroblasts. b. UMAP embedding of the CCA integrated dataset by TRT status. c. UMAP embedding of the CCA integrated dataset by cell type. d. UMAP embedding of the CCA integrated HR+ luminal epithelial cells by data source. e. UMAP embedding of the CCA integrated HR+ luminal epithelial cells by TRT status. f. Dot plot showing decreased expressions of markers of estrogen signaling (PTHLH, DIO2, EREG, AREG, WNT4) in HR+ luminal epithelial cells in donors on TRT, and increased expressions of markers downstream of androgen signaling (RYR2, CUX2, GPC3, SOCS2, NBL1).

**Supplementary Fig. 9.**
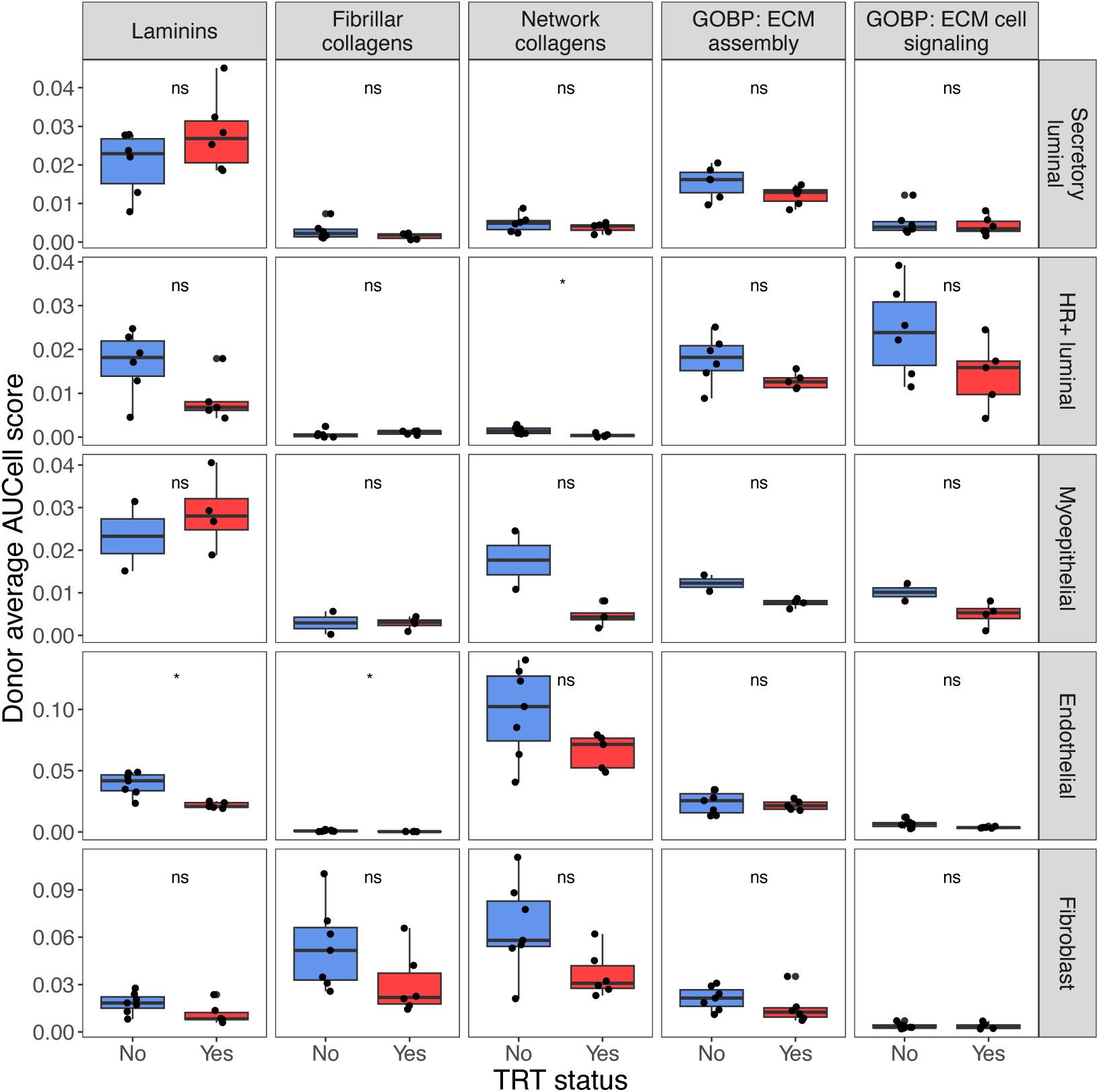
Gene expression comparison of ECM-related genes and GOBP pathways between donors on TRT and donors not on TRT in the current dataset. There is no significant difference in any of the cell types, however, there is a trend in decreased expression in donors on TRT of network collagen genes in myoepithelial cells, endothelial cells, and fibroblasts, as well as a trend for laminins, GOBP ECM assembly, and cell signaling in HR+ luminal cells. However, these results are not statistically significant, suggesting that a larger cohort in future studies should be able to resolve these differences.

**Supplementary table 1:**
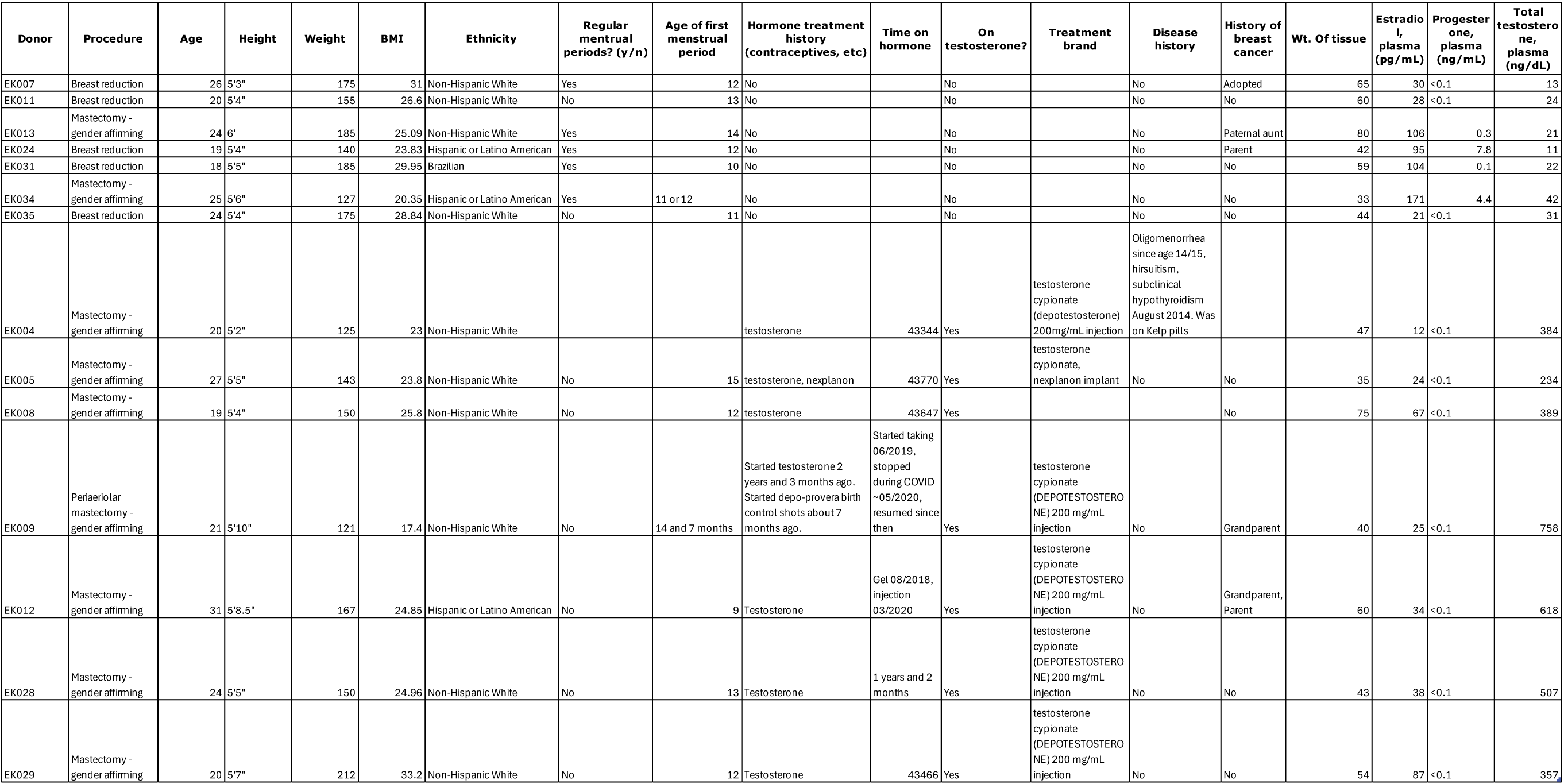
Detailed demographics information for the participants in the single cell RNA sequencing cohort.

**Supplementary table 2:**
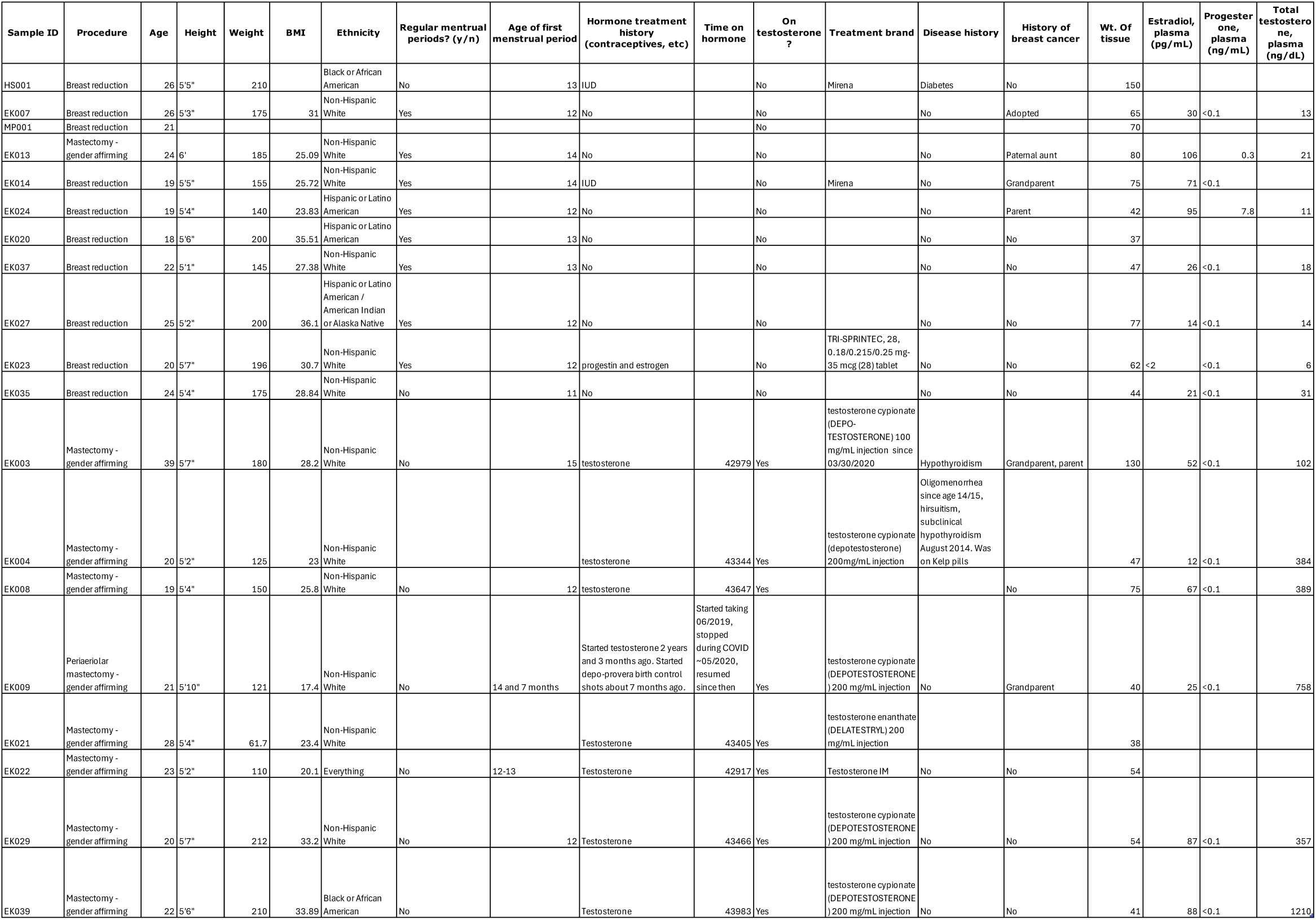
Detailed demographics information for the expanded cohort for immunofluorescence quantifications.

**Supplementary table 3:**
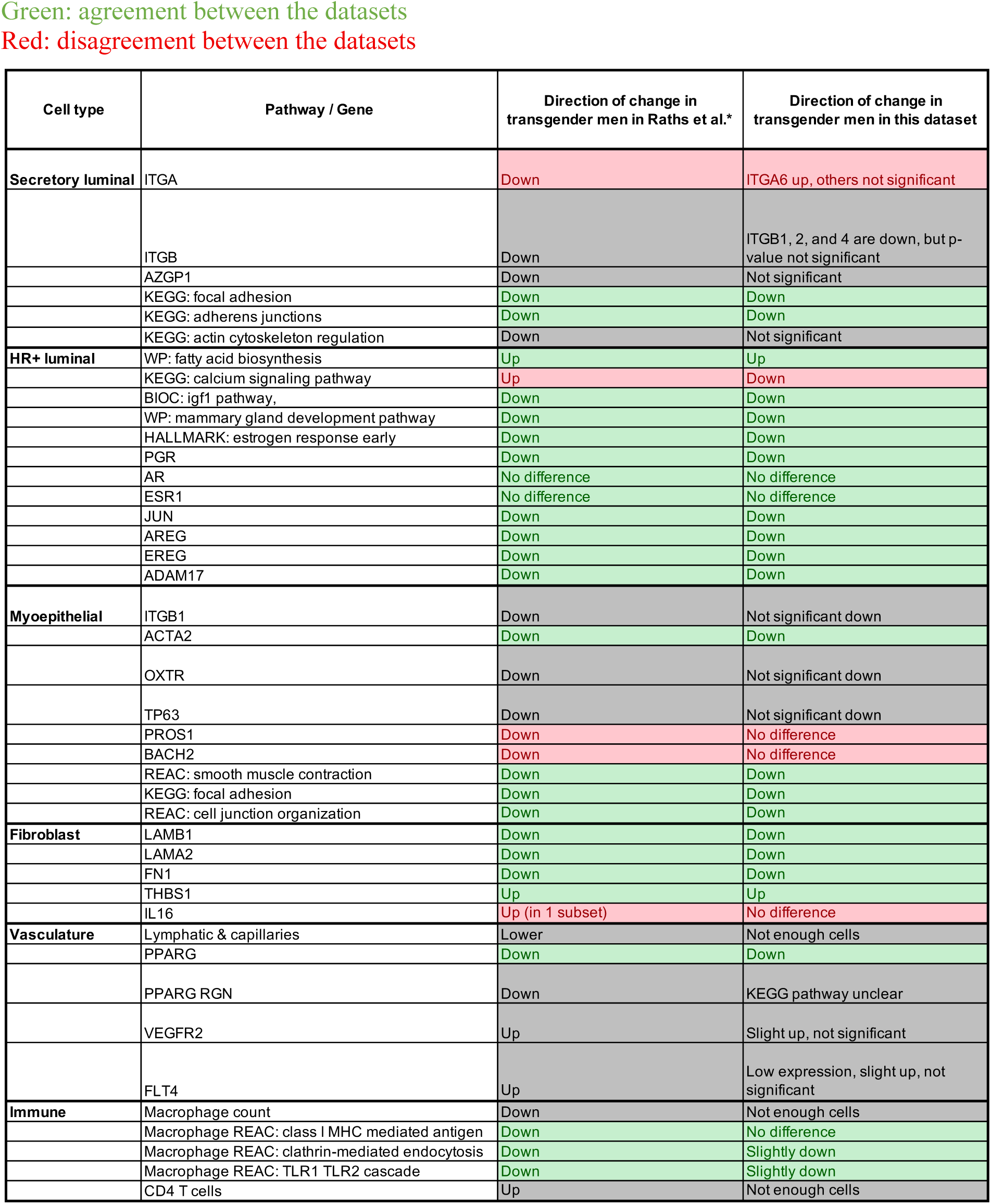
The molecular changes in each cell type in transgender men on testosterone replacement therapy (TRT) relative to cisgender women in the current study, in comparison to a previous study. Data for previous study is derived from Raths F, Karimzadeh M, Ing N, et al (2023) The molecular consequences of androgen activity in the human breast. Cell Genom 3:100272. https://doi.org/10.1016/j.xgen.2023.100272

